# THE ACTIN CYTOSKELETON CONTROLS NADPH OXIDASE ACTIVATION AND G PROTEIN RECRUITMENT MEDIATED BY NEUTROPHIL Gαq-COUPLED RECEPTORS

**DOI:** 10.1101/2025.05.30.656946

**Authors:** Neele K. Levin, Claes Dahlgren, Huamei Forsman, Martina Sundqvist

## Abstract

Signaling by formyl peptide receptor 1 (FPR1), the prototype G protein-coupled receptor (GPCR) expressed in neutrophil leukocytes, is initiated by an activation of a G protein containing a Gα_i_ subunit. FPR1 activation results in an increase in the cytosolic concentration of free calcium ions ([Ca^2+^]_i_), and an activation of the superoxide anion producing NADPH oxidase. Receptor downstream signals generated by the danger molecule ATP recognized by the purinergic receptor P2Y_2_ are transduced by a G protein containing a Gα_q_ subunit. The neutrophil response induced by ATP also includes a transient rise in [Ca^2+^]_i_, but the downstream signals do not activate the NADPH oxidase. ATP can, however, activate this enzyme system through a receptor transactivation mechanism dependent not only on the ATP receptor but also on the free fatty acid receptor FFA2R, provided that this receptor is allosterically modulated. This occurs through a novel mechanism whereby FFA2R is activated from the cytosolic side of the plasma membrane by Gα_q_ transduced signals generated by the ATP receptor. Furthermore, in neutrophils with a disrupted actin cytoskeleton, ATP (as well as platelet activating factor; recognized by the Gα_q_-coupled PAFR) becomes a potent NADPH oxidase activating agonist. At high concentrations of the actin cytoskeleton disrupting drug latrunculin A the activation was only partly reduced by Gα_q_ inhibition. More importantly, this response was also partly inhibited by pertussis toxin. The effects on the ATP-induced NADPH oxidase activity, of the Gα_q_ inhibitor and pertussis toxin were more and less pronounced, respectively, when the concentration of latrunculin A was reduced. Taken together, we show that in primary human neutrophils the actin cytoskeleton is part of the regulatory machinery that determines the activation of NADPH oxidase activation and the G protein recruitment profile downstream of activated of Gα_q_-coupled GPCRs.

**Highlights:**

- ATP is a biased signaling agonist unable to activate the neutrophil NADPH oxidase
- ATP activates the NADPH oxidase through P2Y_2_R mediated transactivation of FFA2R
- Actin cytoskeleton disruption enables ATP to activate the NADPH oxidase
- Cytoskeleton regulated NADPH oxidase activation depends on G_i_ and G_q_ signaling
- The actin cytoskeleton regulates the G protein recruitment profile of P2Y_2_R

## INTRODUCTION

Professional phagocytes such as neutrophil granulocytes and monocytes express an NADPH oxidase with the capacity to transport electrons over a membrane using electrons from NADPH, produced in the hexose monophosphate shunt. The electrons reduce molecular oxygen to superoxide anions (O_2_^-^; [1, 2]). The membrane component of the NADPH oxidase is made up by one larger (NOX2; gp91^phox^) and one smaller protein (p22^phox^) and this heterodimer (cytochrome b_558_) contains two heme groups, and, together with a redox coenzyme (FAD), the cytochrome functions as the catalytic center of the electron transporting enzyme. In resting (naïve) neutrophils several phagocyte oxidase factors (p40^phox^, p47^phox^, p67^phox^, Rac1/2) are cytosolic and for an activation of NADPH oxidase, these proteins must first be recruited to the b cytochrome containing membrane [3]. It has long been well established that signals generated in neutrophils by chemoattractant receptors such as formyl peptide receptor 1 (FPR1), trigger an assembly and activation of the NADPH oxidase in the plasma membrane [4]. As a consequence, the O_2_^-^ produced by neutrophils activated by the FPR1 agonist fMLF are not retained inside the cells but instead released/secreted extracellularly where other reactive oxygen species (ROS) are formed with O_2_^-^ as the starting component (see [5, 6]). Although signaling by activated FPRs has been intensively studied for long, the precise signals and molecular mechanisms that regulate the NADPH oxidase activation process have not yet been identified [7].

Neutrophil receptors and the ligands that regulate their functions constitute the basis for the ability of our innate immune system to recognize both pathogen-associated molecular patterns (PAMPs) and danger-associated molecular patterns (DAMPs). The neutrophil activities induced by such PAMPs and DAMPs are of importance for our ability to combat invading microbes and to repair damaged tissues [8]. Many of the neutrophil plasma membrane-localized receptors belong, just as FPR1, to the family of G protein coupled receptors (GPCRs, also known as seven-transmembrane receptors [7TM receptors] [9]). FPR1 is usually denoted as the prototype for the neutrophil GPCRs and in accordance with name and receptor group affiliation, this receptor is activated by N-formyl methionyl containing peptides and signaling rely on coupling to, and activation of, a heterotrimeric G protein containing a Gα_i_ subunit. Basically, the majority of the initial GPCR activities that are important for receptor downstream signals are mediated by G protein complexes containing a GTP/GDP-binding α subunit with GTPase activity and a dimeric β/γ subunit. In the non-activated and low-signaling state of the receptor, GDP (guanosine diphosphate) is bound to the α subunit, and when the receptor is activated, GTP replaces GDP, and this binding triggers a dissociation of the α subunit from the β/γ subunit. The two dissociated G protein subunits then transfer the information from the activated GPCR to a second messenger cascade specifically characterized by the receptor, the receptor specific ligand, the G protein involved in the response and, the cell in which the receptor is expressed [9, 10]. The dissociated β/γ subunit downstream of FPR1 activates PLC (phospholipase C), an enzyme that catalyzes the hydrolysis of PIP_2_ (phosphatidyl-inositol-4,5-bisphosphate) to IP_3_ (inositol trisphosphate) and DAG (diacylglycerol). The latter is an activator of PKC (protein kinase C). The IP_3_ that is released to the cytoplasm is recognized by IP_3_-receptors on the Ca^2+^ storing endoplasmic reticulum (ER), and when these receptors are occupied, stored Ca^2+^ is released from the ER and the intracellular concentration of free Ca^2+^ ([Ca^2+^]_i_) is increased. Our genome encodes many different G protein subunits, including around 20 α-subunits divided into four groups (Gα_s_, Gα_q_, Gα_i/o_ and Gα_12/13_). These subunits combine with a β/γ dimer generated from the 5 and 12 subunit genes encoding β and γ subunits, respectively [11]. The literature suggests the presence of at least 30 distinct G protein complexes, with the possibility of additional complexes yet to be identified. Studies involving the overexpression of GPCRs in conjunction with various G proteins in cell lines have revealed that certain GPCRs exhibit selective coupling to a limited number of G proteins, while others exhibit promiscuity by coupling to several different G proteins [12]. This promiscuity is expected to increase as more data become available. It is also reasonable to assume that GPCRs previously identified as promiscuous may exhibit a more selective G protein coupling profile when activated in primary cells. The recruitment profile may be contingent not only on the cell in which the receptor is expressed but also on the agonist that activates the receptor [13, 14]. However, the regulatory repertoire is also open to other novel mechanisms that restrict the recruitment of different G proteins. [15]

In addition to FPRs, neutrophils express GPCRs that recognize platelet activating factor (PAF; recognized by PAFR), generated by diverse cells through lipid remodeling, short chain fatty acids (SCFAs, recognized by FFARs) generated by bacteria, and adenosine trisphosphate (ATP; recognized by the purinoreceptor P2Y_2_R) generated by host cell mitochondria [7, 16–19]. Like the FPRs, FFA2R (one of the neutrophil FFARs) couple to a G protein containing a Gα_i_ subunit, and the activated dissociated β/γ subunit activates the PLC-PIP_2_-IP_3_-Ca^2+^ pathway. This signaling pathway is activated also by ATP and PAF but the rise in [Ca^2+^]_i_ downstream of their cognate receptors is mediated by the Gα subunit of a Gα_q_ containing G protein [7]. Despite the signaling similarities, but in contrast to the activation profiles of several neutrophil GPCRs coupled to Gα_i_ containing G proteins, the agonist occupied P2Y_2_R in naïve neutrophils do not generate signals that trigger an assembly and activation of the ROS producing NADPH oxidase [20].

In the context of ATP, we have previously shown that the ATP/P2Y_2_R complex can activate the ROS producing NADPH oxidase by two different mechanisms; i) in TNF-primed neutrophils, ATP triggers such an activation in the presence of an FFA2R selective allosteric modulator, and this activation is achieved through a novel receptor transactivation (cross-talk) mechanism that results in an activation of the allosterically modulated FFA2R [7], and ii) ATP triggers this activation also in neutrophils in which the integrity of the actin cytoskeleton has been disrupted by an actin binding toxin [20]. The precise signaling triggered by ATP leading to an activation of the NADPH oxidase in neutrophils is still poorly understood. We now show that that ATP also activates the NADPH oxidase in neutrophils that have been primed with GM-CSF (granulocyte-macrophage colony-stimulating factor) through a transactivation mechanism involving also FFA2R. The transactivating signals are generated downstream of a Gα_q_ containing G protein. Furthermore, in the context of neutrophils with a disrupted actin cytoskeleton, the activation of the NADPH oxidase by ATP occurs through signals generated both with and without the involvement of a Gα_q_ containing G protein.

## MATERIAL AND METHODS

### Chemicals and reagents

Dextran T500 was from Pharmacosmos and Cytiva Ficoll-Paque^TM^ Plus Medium was from Fischer Scientific. Fura-2-acetoxymethyl ester (AM) was from Invitrogen. Isoluminol, bovine serum albumin (BSA), the actin cytoskeleton disrupter latrunculin A, the Gα_i_ inhibitor pertussis toxin (PTX), the P2Y_2_R agonist ATP, the FFA2R agonist propionic acid (propionate), the FFA2R allosteric modulator Cmp58 ((S)-2-(4-chlorophenyl)-3,3-dimethyl-N-(5-phenylthiazol-2-yl)butanamide), the FPR1 agonist fMLF and the GPR84 agonist ZQ16 were from Sigma-Aldrich. The P2Y_2_R antagonist AR-C118925 ((5-[[5-(2,8-dimethyl-5Hdibenzo[a,d]cyclohepten-5-yl)-3,4-dihydro-2-oxo-4-thioxo-1(2H)-pyrimidinyl]methyl]-N-2H-tetrazol-5-yl-2-furan-carboxamide)) and the FFA2R antagonist CATPB ((S)-3-(2-(3-chlorophenyl)acetamido)-4-(4-(trifluoromethyl)phenyl) butanoic acid) were from Tocris. The PAFR agonist PAF was from Avanti Research. The Gα_q_ inhibitor YM-254890 was acquired from FUJIFILM Wako Pure Chemical Corporation and GM-CSF was purchased from PeproTech. All stock solutions were dissolved in DMSO and subsequent dilutions performed in Krebs-Ringer glucose phosphate buffer (KRG, 120 mM NaCl, 4.9 mM KCl, 1.7 mM KH_2_PO_4_, 8.3 mM Na_2_HPO_4_, 1.5 mM MgSO_4_, 10 mM glucose, and 1 mM CaCl_2_ in dH_2_O, pH 7.3).

### Ethics statement

The present study comprises buffy coats obtained from healthy human blood donors at the blood bank at Sahlgrenska University Hospital in Gothenburg, Sweden. All buffy coats were obtained anonymously and therefore no ethical approval was required according to the Swedish legislation section code 4§ 3p SFS 2003:460 (Law on Ethical Testing of Research Relating to People).

### Isolation of human neutrophils

Neutrophils were isolated from buffy coats of human healthy blood donors as previously described [21, 22] using dextran sedimentation followed by Ficoll-Paque gradient centrifugation. Erythrocytes were lysed by hypotonicity, and the remaining pellet was washed after which the purity was measured on a Sysmex KX-21 N Hematology Analyzer (Sysmex Corporation). All pellets comprised ≥ 90% neutrophils and these were resuspended in KRG to a final neutrophil concentration of 1x10^7^/mL and kept on ice until further analysis the same day. To amplify the activation signals of the NADPH oxidase, the freshly isolated neutrophils were incubated with GM-CSF (2 nM, 20 min, 37°C) at a concentration of 10^6^ neutrophils/mL and then kept on ice until further analysis the same day (these neutrophils are referred to as GM-CSF-primed neutrophils from herein).

### Neutrophil NADPH-oxidase activity

The release of superoxide anions (O_2_^-^) by the neutrophil NADPH oxidase was monitored using an isoluminol-enhanced chemiluminescence (CL) technique as previously described [23, 24]. In short, each 900 μL reaction mix which contained 10^5^/mL neutrophils, 0.2 μM isoluminol and 4 Units/mL HRP was incubated for five min at 37°C prior to the addition of an agonist (100 μL) The NADPH oxidase activity was measured using a six-channel Biolumat LB 9505 (Berthold Co., Wildbad, Germany).

In cases were the effects of certain ligands (inhibitor, antagonist, allosteric modulator, latrunculin A), were studied, these reagents were added either five (inhibitor, antagonist, allosteric modulator) or two minutes (latrunculin A) prior stimulation with the agonist. In experiments were the effect of PTX was studied, GM-CSF-primed neutrophils were incubated without or with PTX (500 ng/mL) for 30 min at 37°C, followed by addition of latrunculin A and stimulation with the agonist.

The O_2_^-^ production was recorded continuously over time and expressed in Mega counts per minute (Mcpm). The peak O_2_^-^ released was used for summary analyses of NADPH oxidase activity.

### Determination of changes in concentration of free calcium ions in the cytosol

Fura-2 AM (2 μM) was added to neutrophils (2 x 10^7^/mL) resuspended in Ca^2+^-free KRG with 0.1% BSA. The cells were incubated in the dark for 30 min followed by washing and resuspension in KRG (2 x 10^7^/mL). The rise in the cytosolic concentration of free Ca^2+^ ([Ca^2+^]_i_) was then measured using a Perkin Elmer fluorescence spectrophotometer (LC50), with excitation wavelengths of 340 nm and 380 nm and emission wavelength of 509 nm. A ratio of the values obtained for 340 nm and 380 nm excited Fura-2 fluorescence over time was calculated and used to express the increase in [Ca^2+^]_i_.

### Data analysis

Data analysis and processing was performed with GraphPad Prism 10.4.1 (GraphPad Software, San Diego, CA, USA). All statistical tests were performed using the peak values of the raw data including the statistically significant differences found in data set where the results are shown as percent of control. The specific statistical test used are described in the figure legends and statistically significant differences are denoted by the following *p*-values: *p* < 0.05 (*), *p* < 0.01 (**), *p* < 0.001 (***) and *p* < 0.0001 (****). Non-statistically significant differences are denoted by ns, i.e., *p* ≥ 0.05.

## RESULTS

### The P2Y_2_R agonist ATP is a functional selective neutrophil activator

In neutrophils, ATP induced a transient elevation in the cytosolic concentration of free Ca^2+^ ([Ca^2+^]_i_). The rise in [Ca^2+^]_i_ was rapidly initiated and reached a peak within the first minute after ATP addition (Fig 1A). This response was inhibited by the specific P2Y_2_R antagonist AR-C118925 (Fig 1A and [25, 26]), consistent with the receptor’s preference for ATP in neutrophils. P2Y_2_R belongs to the family of G protein-coupled receptors (GPCRs), and the [Ca^2+^]_i_ rise induced by GPCRs may be attributed to an activated Gα_q_ subunit as well as a βγ dimer separated from its Gα_i_ subunit in the receptor-coupled G protein [27, 28]. The inhibitory effect of the Gα_q_ inhibitor YM-254890 demonstrated that the ATP-activated purine receptor P2Y_2_R couples to a Gα_q_-containing G protein (Fig 1A). In contrast, the transient rise in [Ca^2+^]_i_ induced by the formylated peptide fMLF, specifically recognized by the Gα_i_-coupled FPR1, was unaffected by YM-254890 (Fig 1B).

**Figure 1.**
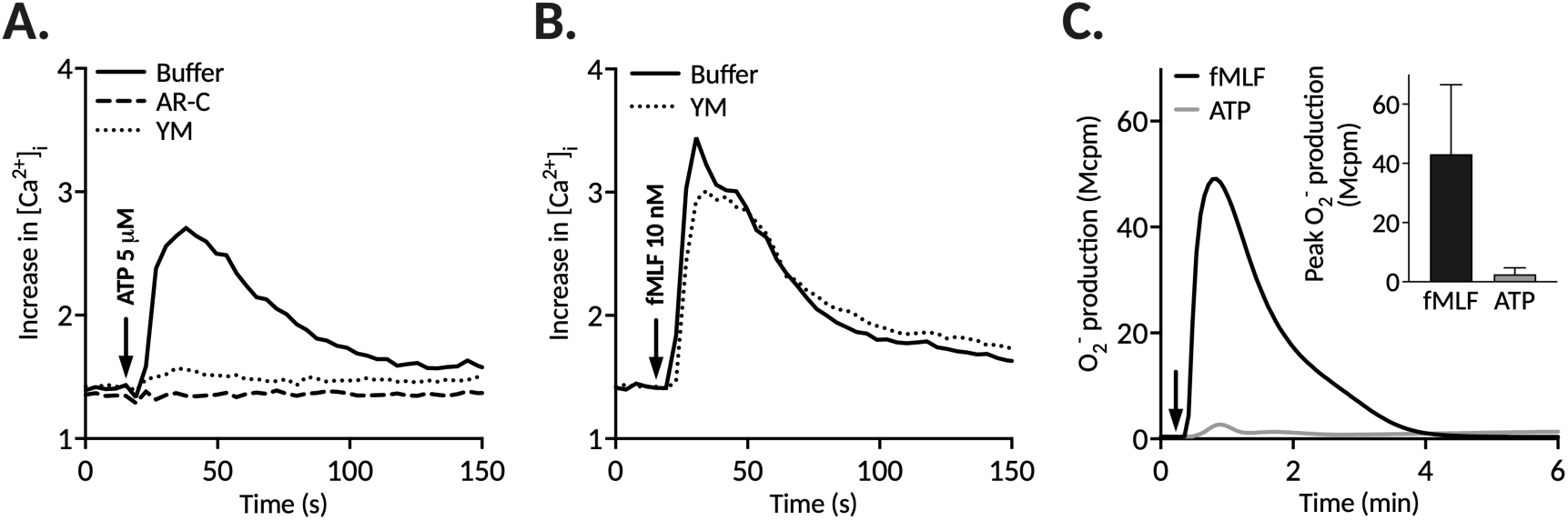
ATP triggers a transient increase in intracellular Ca²⁺ ([Ca²⁺]_i_) but does not activate the NADPH oxidase in human neutrophils. **A-B.** Fura-2-loaded neutrophils were pre-incubated (10 min, 37°C) with either buffer, the P2Y₂R-specific antagonist AR-C118925 (AR-C, 1 μM), or the Gα_q_ inhibitor YM-254890 (YM, 200 nM). Thereafter **A.** ATP (5 μM) or **B.** fMLF (10 nM) was added (arrow indicates time of addition) and the transient rise in [Ca²⁺]_i_, was measured over time. Representative traces from an individual experiment are shown. **C**. Superoxide anion (O_2_^-^) production measured in GM-CSF-primed neutrophils stimulated with ATP (50 μM) or fMLF (25 nM) over time (arrow indicates time of addition). **Inset**. The peak O₂^⁻^ production after stimulation with ATP (n=18) or fMLF (n=4) are summarized in the bar graph (mean ± SD).

The binding of fMLF to its neutrophil receptor, FPR1, was associated with an assembly of the NADPH oxidase and by that a generation of superoxide anions (Fig 1C). In contrast, ATP did not induce this activation (Fig 1C), consistent with previous findings that there is no direct correlation between an increase in [Ca^2+^]_i_ and NADPH oxidase activation [7].

### GM-CSF primes the NADPH oxidase activity in human neutrophils

In accordance with earlier findings [29], the NADPH oxidase activity triggered by fMLF in neutrophils preincubated with GM-CSF was primed, i.e., the response was substantially increased compared to the response induced in naïve neutrophils (Fig 2).

**Figure 2.**
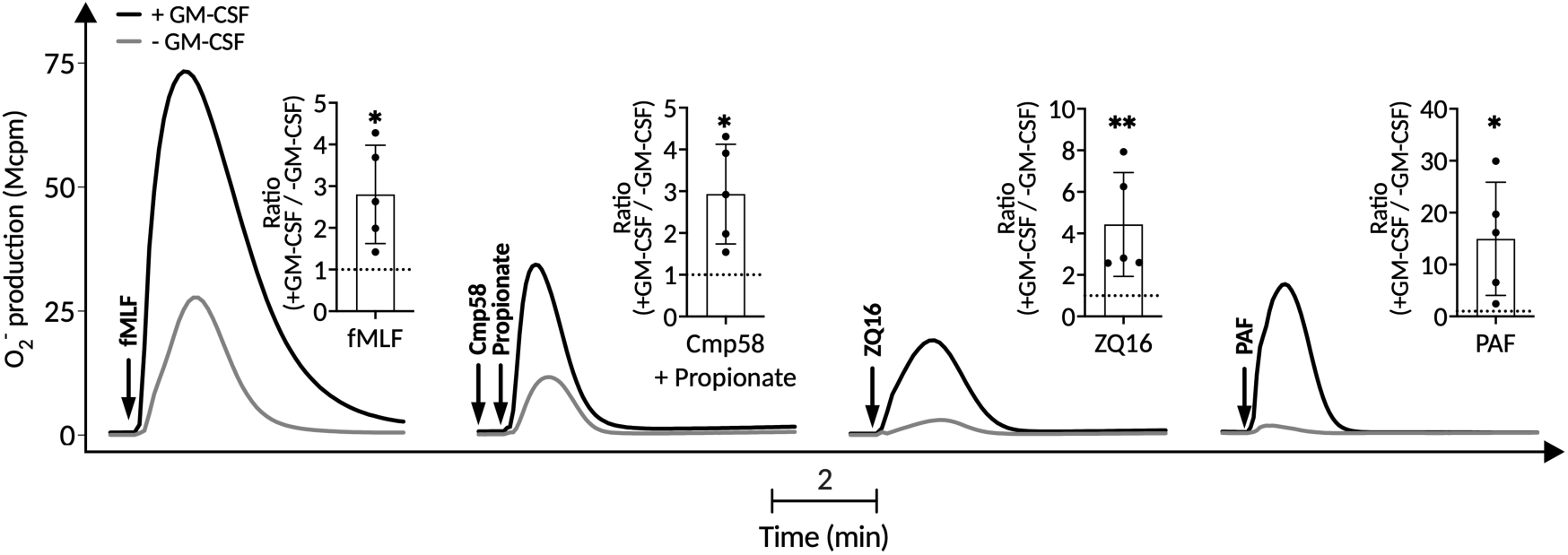
GM-CSF primes neutrophils -- the NADPH oxidase activity in response to various GPCR agonists is increased. Neutrophils incubated with or without GM-CSF (2 nM, 20 min, 37°C) were activated with either the FPR1 agonist fMLF (25 nM), the FFA2R agonist propionate (25 µM, combined with in the allosteric FFA2R modulator Cmp58, 1 µM), the GPR84 agonist ZQ16 (1 μM), or the PAFR agonist PAF (100 nM) and the O_2_^-^ production was monitored over time. Representative traces for each agonist are shown. **Insets**. The ratio of the peak O_2_^-^ production in GM-CSF-primed neutrophils (+ GM-CSF) versus unprimed cells (-GM-CSF) are shown (mean ± SD, n=5). The dotted horizontal line in each bar graph represents a ratio of 1. Statistically significant differences between GM-CSF-primed and unprimed neutrophils were evaluated by a paired Student’s *t*-test for each agonist separately (\**p* < 0.05, \*\**p* < 0.01).

The ROS production was also augmented in neutrophils pre-treated with GM-CSF compared to naïve neutrophils stimulated with either propionate (FFA2R agonist) combined with the allosteric FFA2R modulator Cmp58, ZQ16 (GPR84 agonist), or PAF (a PAFR agonist) (Fig. 2). Like the inability for ATP to activate the NADPH oxidase in naïve neutrophils, ATP on its own did not trigger any NADPH oxidase activity in GM-CSF-primed neutrophils (Fig 3).

**Figure 3.**
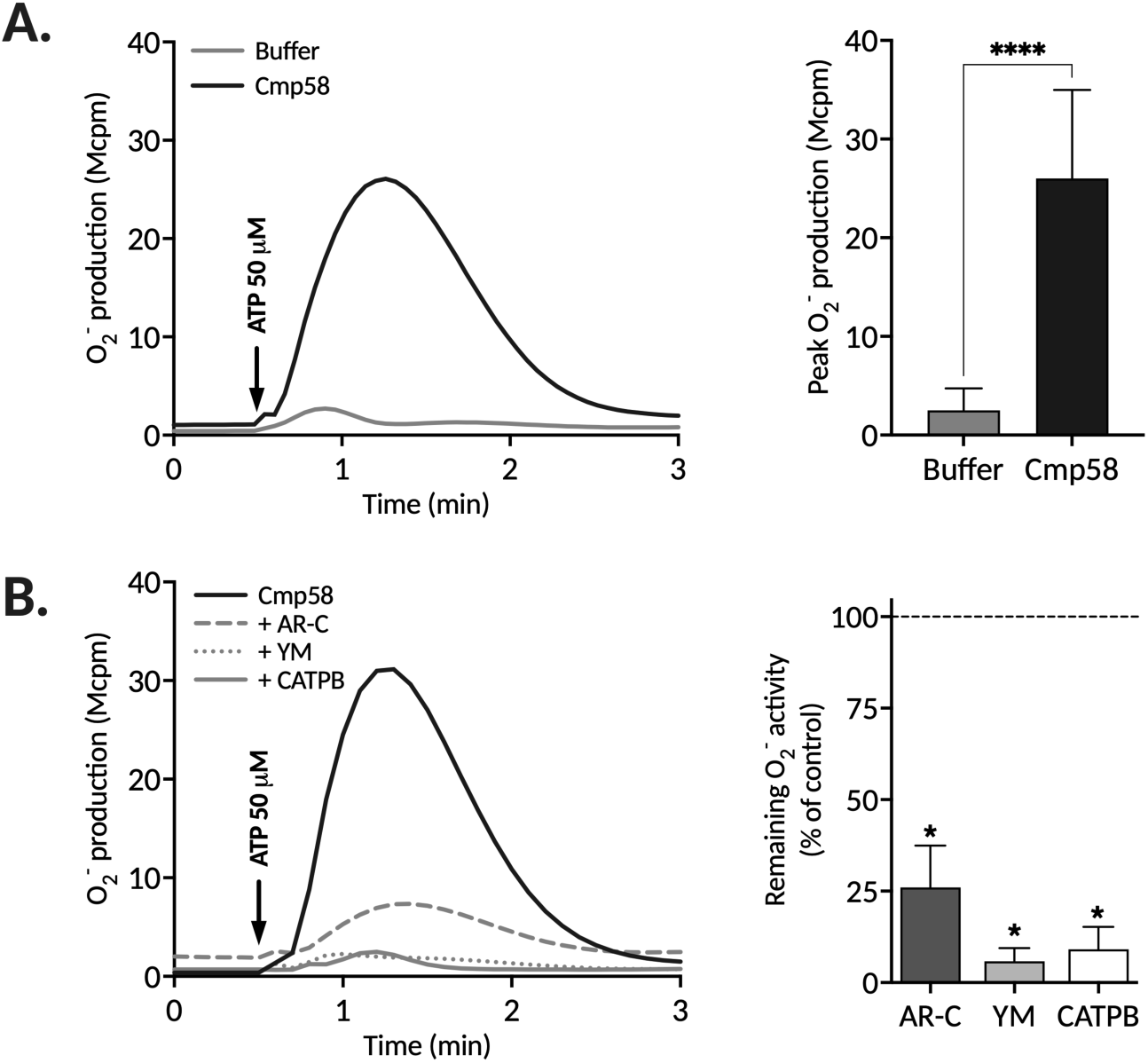
Allosteric modulation of FFA2R converts ATP into an activator of the neutrophil NADPH oxidase. The O_2_^-^ production was measured in GM-CSF-primed neutrophils pre-incubated (5 min, 37°C) with either buffer, the allosteric FFA2R modulator Cmp58 (1 μM), the Gα_q_ inhibitor YM-254890 (YM, 200 nM), the P2Y₂R-specific antagonist AR-C118925 (AR-C, 1 μM), or the FFA2R-specific antagonist CATPB (100 nM) prior stimulation with ATP (50 μM; arrows indicate time of addition). **A.** Left: A representative trace of the O_2_^-^ production induced by ATP in the absence or presence of Cmp58 is shown. Right: Summary of the peak O₂^⁻^ production (mean ± SD, n=18). **B.** Left: A representative trace of the O_2_^-^ production induced by ATP in the presence of Cmp58 alone or in combination with YM, AR-C or CATPB is shown. Right: Summary of the peak O_2_^-^ production (mean ± SD, n=5). Statistically significant differences in **A**. were evaluated by a paired Student’s *t*-test and in **B**. with a repeated measures one-way ANOVA followed by Dunnett’s multiple comparisons test to Cmp58 alone control (\**p* < 0.05, \*\*\*\**p* < 0.0001).

Taken together, as compared to untreated (naïve) neutrophils, the cytokine GM-CSF-primed neutrophils to induce increased ROS production in response to several GPCR agonists. Based on these results, GM-CSF priming was used to amplify the NAPDH oxidase activation signals in response to GPCR agonists for the rest of the experiments in this study.

#### Allosteric modulation of FFA2R turns ATP into an NADPH oxidase activating agonist

As mentioned, ATP on its own did not trigger any NADPH oxidase activity in GM-CSF primed neutrophils (Fig 3A). However, the inability of the signals from P2Y_2_R to activate the neutrophil NADPH oxidase was not absolute. In the presence of the positive allosteric FFA2R modulator Cmp58, ATP induced a release of superoxide anions (O_2_^-^) from neutrophils (Fig 3A). This ATP-induced response was blocked by the specific FFA2R antagonist CATPB, confirming the involvement of FFA2R (Fig 3B). As anticipated, the response was also inhibited by the P2Y_2_R-specific antagonist AR-C118925 and the Gα_q_-selective inhibitor YM-254890 (Fig 3B). Our model for how this activation is achieved assumes that Gα_q_ dependent signals induced by ATP transactivates FFA2R from the cytosolic side of the plasma membrane and the transactivated receptor subsequently activate the NADPH oxidase.

### Latrunculin A turns ATP into an activator of the NADPH oxidase and concentration-dependently renders the response insensitive to Gα_q_-inhibition

Latrunculin A is a research tool-toxin that binds actin monomers and prevents them from participating in the polymerization process; binding of the toxin, thus, results in a disruption of the actin filaments of the cell cytoskeleton [30]. When Cmp58 was replaced by latrunculin A, ATP was converted into an NADPH oxidase activating agonist and the effect of a depolymerized actin cytoskeleton was clearly more dramatic (Fig 4). At latrunculin A concentrations above 25 ng/mL, there was essentially no further increase in the ATP-induced NADPH oxidase activity (data not shown). The NADPH oxidase activity was dependent on the concentration of ATP and the effect was most pronounced with a latrunculin A concentration of 25 ng/mL (Fig 4). Based on the inhibitory effect of the Gα_q_-selective inhibitor YM-254890 on the downstream signaling of the activated P2Y_2_R, we assumed that the inhibitor would potently reduce the response induced by ATP in the presence of latrunculin A. However, our data show that the inhibition mediated by YM-254890, was not very pronounced when a high concentration of latrunculin A was added to the system (Fig 4A). This lack of inhibition was evident irrespectively of the ATP concentration used to activate the cells. With ATP concentrations of 50 µM and 1 µM the inhibition was around 25% and 50%, respectively (Fig 4A). At latrunculin A concentrations below 25 ng/mL, the level of NADPH oxidase activity was gradually reduced in relation to the concentration of the toxin (Fig 4B and C). In contrast, the inhibitory effects of the Gα_q_-selective inhibitor YM-254890 on the ATP induced activation of the NADPH oxidase, was gradually increased in relation to the concentration of latrunculin A, reaching a level of around 90% with a latrunculin A concentration of 5 ng/mL (Fig 4C).

**Figure 4.**
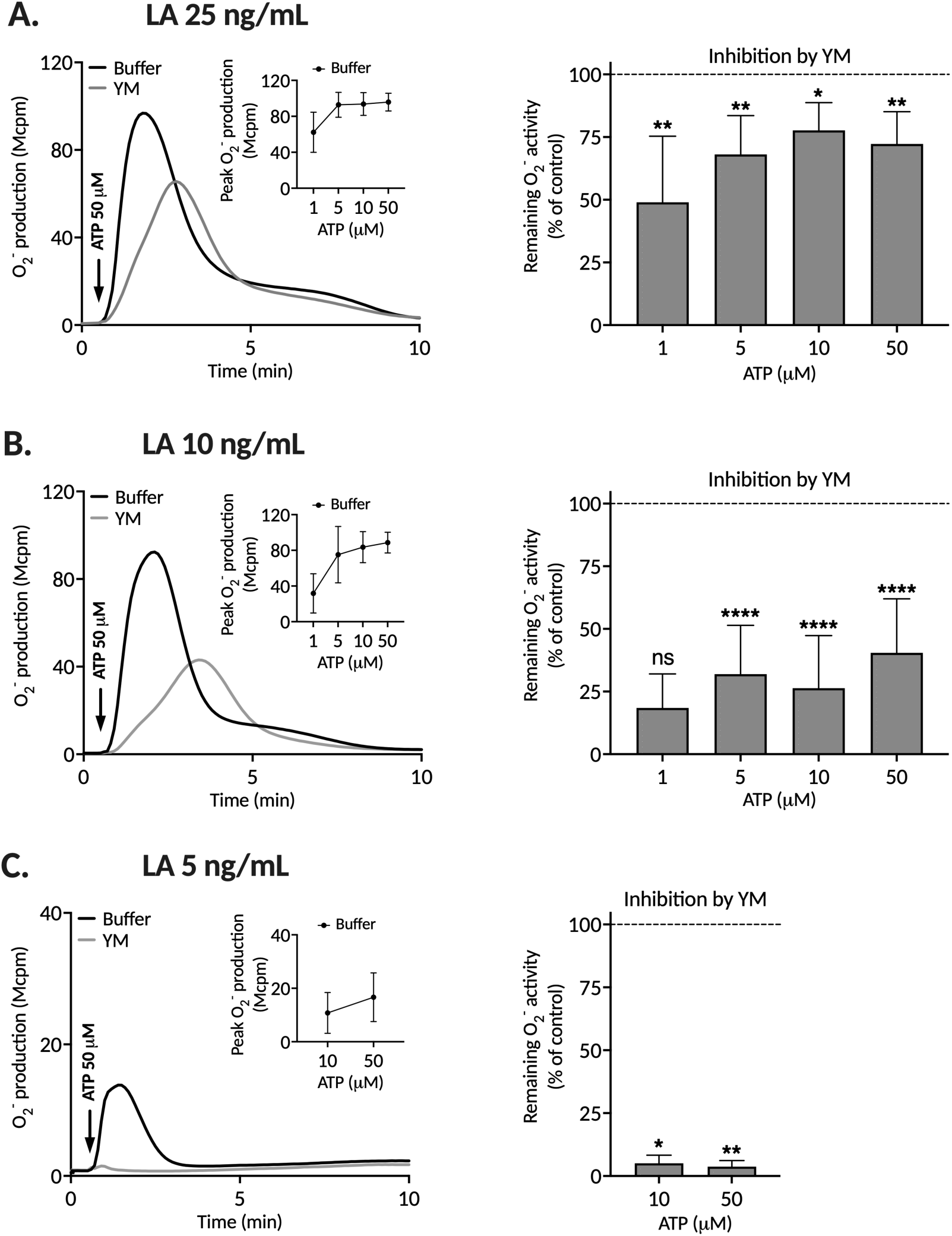
Latrunculin A converts ATP into an activator of the neutrophil NADPH oxidase and changes the sensitivity of the response to Gα_q_-inhibition. ATP induced superoxide anion (O_2_^-^) production measured in GM-CSF-primed neutrophils pre-incubated with different concentrations of latrunculin A (LA) in the absence or presence of the Gα_q_ inhibitor YM-254890 (YM, 200 nM). **A.** Left: Representative traces of the O_2_^-^ production induced by ATP (50 μM; arrow indicates time of addition) in the presence of 25 ng/mL LA with or without YM. Inset: The graph displays peak O_2_^-^ values triggered by different concentrations of ATP (1, 5, 10, or 50 μM) in the presence of 25 ng/mL LA. Right: The bar graph shows the remaining O_2_^-^ activity after ATP (1, 5, 10, or 50 μM) activation of neutrophils in the presence of YM and LA (25 ng/mL). **B–C.** The same experimental setup as in panel A, but with 10 ng/mL (**B**) or 5 ng/mL (**C)** LA, respectively. Insets and bar graphs are represented as mean ± SD (n=5). Statistically significant differences in bar graphs were evaluated by repeated measures one-way ANOVA followed by Šídák’s multiple comparisons test for analysis of each ATP concentration in the absence or presence of YM, separately (\**p* < 0.05, \*\**p* < 0.01, \*\*\*\**p* < 0.0001, ns = not significant).

Taken together, these data show that ATP has the capacity to activate the neutrophil NADPH oxidase by signals generated downstream of a Gα_q-_containing G protein as well as signals generated independent of such a G protein.

#### Latrunculin A affects the sensitivity to Gα_q_ inhibition also in neutrophils activated by PAF

A comparison of the neutrophil-expressed GPCRs that recognize ATP and PAF reveals both similarities and differences [7]. One difference being that PAF is an activator of the NADPH oxidase also in neutrophils with an intact actin cytoskeleton. A similarity is that they both couple to a Gα_q_ containing G protein, and in accordance with this, the PAF-induced response was sensitive to Gα_q_ inhibition (Fig 5A). Regarding the impact of a disruption of the actin cytoskeleton, a similar pattern as that described for the ATP induced response (Fig 4), also characterized the response induced by PAF. That is, the NADPH oxidase activity was substantially increased in the presence of latrunculin A and at high concentrations of the toxin (10 and 25 ng/mL, respectively) the response was only partly reduced by the Gα_q_ inhibitor (Fig 5B and C). However, when the latrunculin A concentration was reduced to 5 ng/mL, the PAF-induced response was once again dependent on Gα_q_-signaling as the Gα_q_ inhibitor reduced the response by ≈ 95% (Fig 5D). In summary, these data show that not only ATP but also PAF, has the capacity to activate the neutrophil NADPH oxidase via both Gα_q_-dependent and Gα_q_-independent signals.

**Figure 5.**
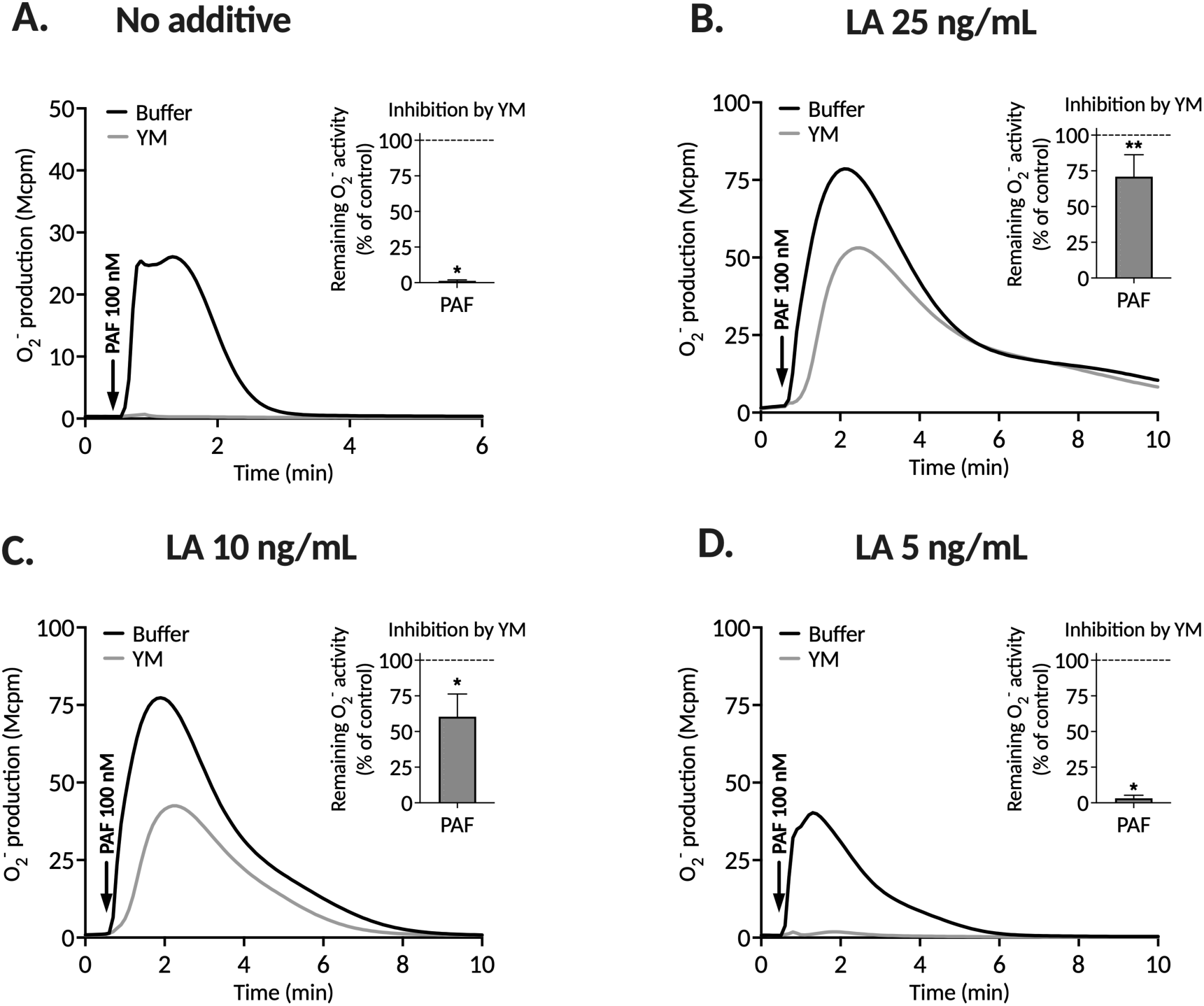
Latrunculin A potentiates the PAF-induced neutrophil NADPH oxidase activity and changes the sensitivity of the response to Gα_q_-inhibition. PAF induced O_2_^-^ production measured in GM-CSF-primed neutrophils incubated without or with different concentrations of latrunculin A (LA) in the absence or presence of the Gα_q_ inhibitor YM-254890 (YM, 200 nM) prior stimulation with PAF (100 nM; arrow indicates time of addition). **A.** A representative trace of the neutrophil O_2_^-^ production when activated by PAF (100nM, arrow indicates time of addition) without any LA and in the absence or presence of YM. **Inset**: The bar graph shows the remaining NADPH oxidase activity induced in neutrophils PAF in the presence of YM. **B–D.** The same experimental setup as in panel A, but with the PAF-induced NADPH oxidase activity in the presence of 25 ng/mL (**B**), 10 ng/mL (**C**) or 5 ng/mL (**D**) LA. **Insets**: Inhibition of the response by YM expressed as remaining activity in the presence of the inhibitor (mean ± SD, n=5). Statistically significant differences in the bar graphs were evaluated by a paired Student’s *t*-test (\**p* < 0.05, \*\**p* < 0.01).

#### Pertussis toxin sensitivity reveals Gα_i_ involvement in the ATP induced NADPH oxidase activation

To further elucidate the intracellular signaling pathway in Gα_q_-independent ATP induced activation of the NADPH oxidase, we employed pertussis toxin (PTX) which, although the G protein subtype selectivity of PTX has been questioned [7], is an established inhibitor of signaling by GPCRs that couple to Gα_i_-containing G proteins [31]. Neutrophils treated with latrunculin A at a concentration of 25 ng/mL and 10 ng/mL, exhibited approximately 40% and 60% remaining ATP response (relative to control cells not treated with PTX, Fig 6A and B). Conversely, PTX lacked inhibitory effect on the ATP induced neutrophil response when the latrunculin A concentration was reduced to 5 ng/mL (Fig 6C). Taken together, the data presented clearly show that in neutrophils treated with high concentrations of latrunculin A, the downstream signals that activates the NADPH oxidase utilizes signals that depend on Gα_q_ and a switch to a Gα_i_ initiated pathway. Conversely, no G protein switch occurs when the concentration of latrunculin A is reduced, yet signaling proceeding exclusively via Gα_q_ still activates the NADPH oxidase.

**Figure 6.**
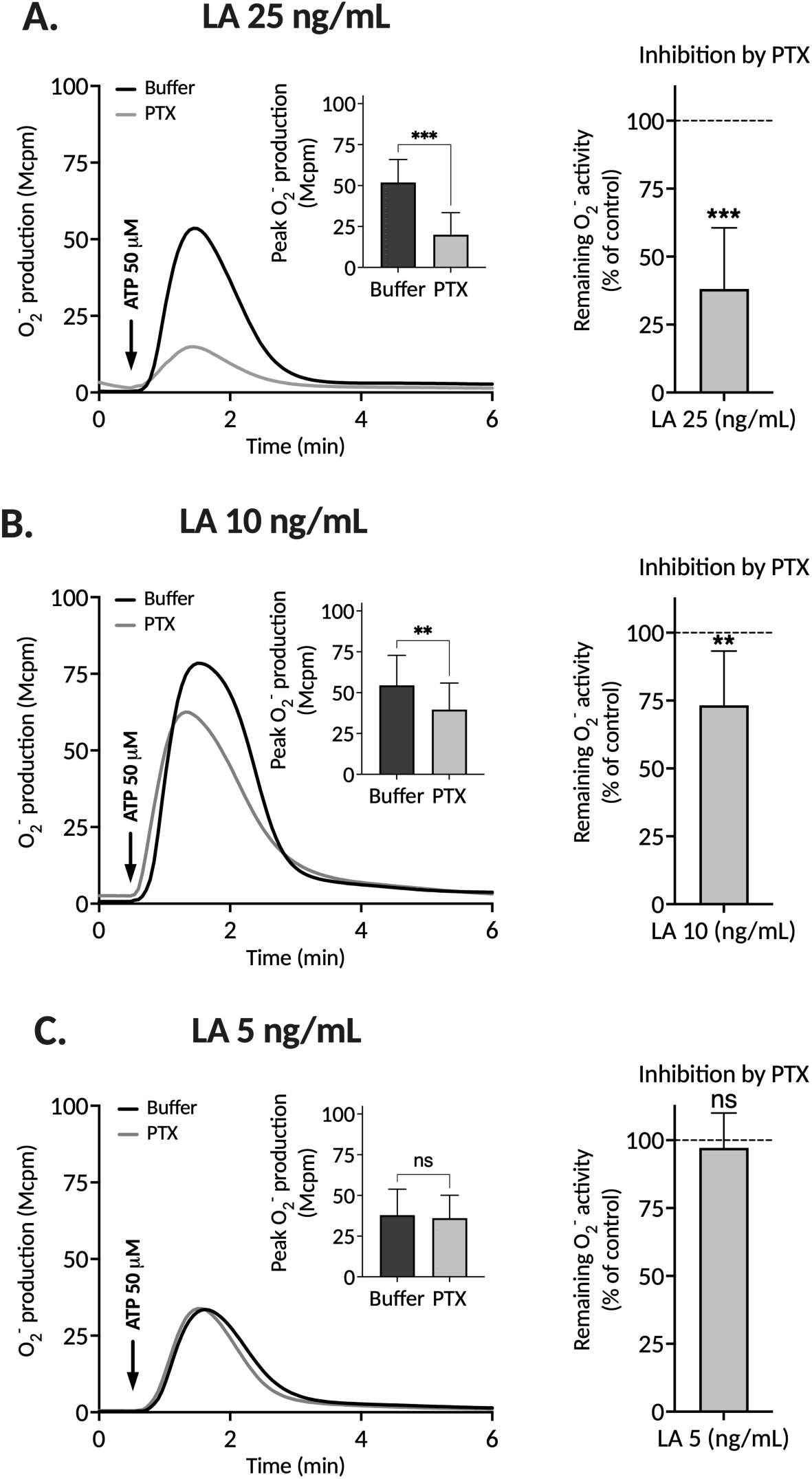
The ATP-induced neutrophil NADPH oxidase activity is sensitive to pertussis toxin in the presence of high latrunculin A concentrations. Superoxide anion (O_2_^-^) production was measured in GM-CSF-primed neutrophils, incubated without (control) or with pertussis toxin (PTX, 500 ng/mL; 30 min at 37°C). These neutrophils were activated by ATP (50µM; arrow indicates time of addition) in the presence of different concentrations LA. **A. Left:** Representative traces of the O_2_^-^ production in PTX treated and control neutrophils activated by ATP (50µM) in the presence of 25 ng/mL latrunculin A (LA). **Inset:** Peak values of the ATP induced NADPH oxidase activity in PTX treated and control neutrophils (mean±SD, n=7). **Right:** Inhibitory effect of PTX, expressed as remaining NADPH oxidase activity induced by ATP in neutrophils treated with PTX. **B–C.** The same experimental setup as in panel A, but with 10 ng/mL (**B**) or 5 ng/mL (**C**) LA, respectively. Statistically significant differences in the bar graphs were evaluated by a paired Student’s *t*-test (\*\**p* < 0.01, \*\*\**p* < 0.001, ns = not significant).

## DISCUSSION

The receptor downstream signals generated in neutrophils by the agonist/receptor pair ATP/P2Y_2_R fulfills the basic characteristic of an agonist that through binding to its receptor induces a “functional selective” response that is the result of a “biased receptor signaling cascade” [32]. This means that in contrast to a balanced GPCR agonists such as the FPR1 agonist fMLF, which facilitates signaling that regulates several different neutrophil functions, the activated ATP receptor transfers signals that result in a functional profile that is circumcised. Agonist binding both to P2Y_2_R and FPR1 results in coupling of the two receptors to G proteins, which activate phospholipase C and this activation gives directly rise to a production of inositol trisphosphate (IP_3_), which is linked to an increase in the cytosolic concentration of free calcium ions ([Ca^2+^]_i_,) [20, 33]. Although this response is induced by both receptors it is transduced by different G proteins; whereas FPR1 couples to a Gα_i_ containing G protein, the signaling G protein partner of P2Y_2_R contains an α_q_ subunit (this study and [7]). Despite the signaling similarity, the functional outcome in naïve neutrophils differs. The signals that trigger an assembly and activation of the superoxide anion generating NADPH oxidase are not generated by the ATP activated P2Y_2_R, clearly showing that the downstream signals generated are not alone sufficient to activate the oxidase. The incapacity of ATP to activate the NADPH oxidase may be due either to that no NADPH oxidase activating signals can be produced by the Gα_q_-coupled P2Y_2_R, or that such signals can be produced also by this receptor and its G protein partner, but that this signaling pathway is (in one way or another) blocked in naïve neutrophils. The primary signaling mechanism downstream of the activated P2Y_2_R involves coupling to the Gα_q_-containing G protein and this signaling synergize with other neutrophil receptors such as the free fatty acid 2 receptor (FFA2R). Our data show that in the presence of an FFA2R specific positive allosteric modulator, the P2Y_2_R agonist ATP activates the neutrophil NADPH oxidase. We have earlier shown that PAF shares the basic primary signaling mechanism with ATP, involving coupling to Gα_q_ and activation of the neutrophil NADPH oxidase by signals that synergize with FFA2R in the presence of a positive allosteric FFA2R modulator [34].

The actin-rich cortex of neutrophils is, together with several different actin binding proteins, closely associated with the cytosolic side of the plasma membrane. It can therefore be deduced that an interaction between cytoskeleton proteins and integral plasma membrane proteins such as GPCRs may affect the function of the latter. In neutrophils, this has primarily been studied with FPR1 as a model for GPCRs. The basic regulatory function of the actin cytoskeleton in FPR1 signaling was described over 30 years ago in some excellent publications by Algirdas Jesaitis and co-workers [35–39]. Little notice has been given to these early observations, which have been confirmed and extended to also include the closely related FPR2 [7, 15]. Based on the results obtained in these studies, a working model that describes how FPR signaling is regulated has been developed; signaling by FPRs is terminated by actin cytoskeleton dependent receptor desensitization, a process in which a continuous G protein recruitment to the activated FPRs becomes physically blocked when the actin cytoskeleton binds to the activated receptor. The molecular mechanism through which the cytoskeletal proteins bind to the cytosolic parts of the FPRs remains to be elucidated, but the results obtained in studies of the neutrophil NADPH oxidase activity, induced by FPR agonists in the presence of inhibitors of actin polymerization, such as latrunculin A, provide strong support for the suggested model [7, 15]. However, a somewhat divergent regulatory profile emerges when the model is applied to analyze the data obtained when ATP is used as the activating agonist instead of FPR agonists. In naïve neutrophils, the NADPH oxidase activating signals are evidently not generated by the activated ATP receptor unless the actin cytoskeleton first has been disrupted. This suggests that in neutrophils, the signaling pathway leading to an activation of the NADPH oxidase is blocked by the cytoskeleton whereas the pathway resulting in a transient rise in [Ca^2+^]_i_ is accessible. Furthermore, the prolongation of the response, characterizing the response induced by FPR agonists in neutrophils with a disrupted actin cytoskeleton [40–42], is absent in the response to ATP, suggesting that this response is terminated without the involvement of the actin cytoskeleton. More importantly, however, that the inhibitory effect of the Gα_q_ inhibitor diminishes with increasing concentrations of latrunculin A. These data suggest on the one hand, that NADPH oxidase activating signals are generated through an involvement of a Gα_q_-containing G protein and, on the other hand, that such signals are generated also by Gα_q_-independent signals. In agreement with this, the ATP induced response in the presence of high latrunculin A concentrations is partly reduced by pertussis toxin, an inhibitor of Gα_i_-mediated signaling. Conversely, the ATP induced response in the presence of low latrunculin A concentrations is entirely dependent on Gα_q_, thereby evading inhibition by PTX. The effect of a disruption of the actin cytoskeleton on the response induced by PAF, an agonist recognized by a receptor (PAFR) that, like P2Y_2_R, couples to a Gα_q_ containing G protein, is analogous to that described for the ATP-induced response, thereby adding another functional similarity for these two neutrophil-expressed GPCRs.

### Conclusions

A disruption of the actin cytoskeleton affects G protein involvement in ATP-induced signals that activate the neutrophil NADPH oxidase; the Gα_q_ signals that activate the NADPH oxidase are apparently blocked in neutrophils with an intact actin cytoskeleton, whereas such signals are generated when this block is removed by the actin cytoskeleton disrupting drug latrunculin A. In addition, a disrupted cytoskeleton allows for a change of the G protein recruitment pattern. As such, also Gα_i_ dependent signals activate the NADPH oxidase in ATP activated neutrophils with a disrupted cytoskeleton. The G protein coupling maps for a multitude of GPCRs reveal that, whereas some GPCRs selectively activate a limited number of Gα proteins, others have the capacity to couple promiscuously to a large number of different Gα subtypes [12]. The purinergic receptor P2Y_2_R belongs to the group of promiscuous GPCRs, capable of recruiting multiple G proteins, whereas the PAFR displays a more restricted coupling profile that has been shown not to engage Gα_q_ [12]. Nevertheless, it is evident from the data presented, that in neutrophils the immediate signals generated, when ATP as well as PAF bind to their respective receptor, are primarily mediated by a Gα_q_-containing G protein. This raises questions concerning the determinants of such a single-faceted selectivity profile. The molecular basis for the G protein selectivity could of course be related to the fact that cells/tissues differ in their G protein expression profile, but other mechanisms may regulate the G protein recruitment and signaling processes, mechanisms that allow responding cells to alter intracellular signaling pathways and the functional outcome of an agonist-induced activation. It has been shown that binding of an allosteric modulator to the prostaglandin 2 receptor EP4, occupying a receptor site localized close to the binding site for the recruited G protein complex, affects the G protein recruitment profile. That is, binding of a positive allosteric modulator to an intracellular receptor site in EP4, converts the signaling from being Gα_s_-dependent to instead use Gα_i_ as the signaling partner [43]. However, G protein selectivity most likely relies on a multitude of other molecular mechanisms, including structural conformation alterations in intracellular G protein binding receptor interfaces, induced from within the cell by regulatory proteins that dynamically facilitate or selectively block G protein binding. Collectively, such mechanisms ensure that GPCRs engage the appropriate G protein signaling pathway in response to different stimuli and in neutrophils the actin cytoskeleton is suggested to be part of this regulatory machinery. It is reasonable to assume that other binding partners present on the cytosolic side of the receptor-expressing plasma membrane may also have the capacity to be unbound or bound to a receptor, thereby regulating GPCR signaling through an interaction with the parts of a receptor that are accessible from the cytosolic side of the plasma membrane. Although the precise mechanisms by which the GPCR mediated recruitment of G proteins is regulated in neutrophils are unknown, it is evident that the actin cytoskeleton has a regulatory role that is crucial for the signaling and activation profile in ATP-activated neutrophils.

## Author contributions

**Neele K. Levin:** Conducted experiments, analyzed and interpreted data, and designed new experiments; Writing - review and editing.

**Claes Dahlgren**: Supervised, analyzed and interpreted data, and designed new experiments; Writing - original draft preparation.

**Huamei Forsman**: Supervised, analyzed and interpreted data, and designed new experiments; Writing - review and editing.

**Martina Sundqvist**: Conceptualization - original idea; Supervised, analyzed and interpreted data, and designed new experiments; Writing - review and editing.

## Abbreviations

ATP: adenosine triphosphate
CL: chemiluminescence
DAMP: danger associated molecular pattern
ER: endoplasmic reticulum
fMLF: formyl-Met-Leu-Phe
FPR: formyl peptide receptor
FFA2R: free fatty acid receptor 2
GDP: guanosine diphosphate
GM-CSF: granulocyte-macrophage colony-stimulating factor
GPCR: G protein-coupled receptor
GTP: guanosine triphosphate
HRP: horseradish peroxidase
IP_3_: inositol trisphosphate
KRG: Krebs-Ringer glucose
LA: latrunculin A
PAF: platelet-activating factor
PAMP: pathogen associated molecular pattern

## REFERENCES

[1] W.M. Nauseef, The phagocyte NOX2 NADPH oxidase in microbial killing and cell signaling, Curr Opin Immunol 60 (2019) 130–140.

[2] E. Pick, NADPH-oxidases revisited:from function to structure, Springer Nature, Cham, Switzerland, 2023.

[3] W.M. Nauseef, R.A. Clark, Intersecting Stories of the Phagocyte NADPH Oxidase and Chronic Granulomatous Disease, Methods Mol Biol 1982 (2019) 3–16.

[4] R.D. Ye, F. Boulay, J.M. Wang, C. Dahlgren, C. Gerard, M. Parmentier, C.N. Serhan, P.M. Murphy, International Union of Basic and Clinical Pharmacology. LXXIII. Nomenclature for the formyl peptide receptor (FPR) family, Pharmacol Rev 61(2) (2009) 119–61.

[5] J. Bylund, K.L. Brown, C. Movitz, C. Dahlgren, A. Karlsson, Intracellular generation of superoxide by the phagocyte NADPH oxidase: how, where, and what for?, Free Radic Biol Med 49(12) (2010) 1834–45.

[6] C. Dahlgren, A. Karlsson, J. Bylund, Intracellular Neutrophil Oxidants: From Laboratory Curiosity to Clinical Reality, J Immunol 202(11) (2019) 3127–3134.

[7] C. Dahlgren, H. Forsman, M. Sundqvist, L. Bjorkman, J. Martensson, Signaling by Neutrophil G Protein-Coupled Receptors that Regulate the Release of Superoxide Anions, J Leukoc Biol (2024).

[8] M. Metzemaekers, B. Malengier-Devlies, M. Gouwy, L. De Somer, F.Q. Cunha, G. Opdenakker, P. Proost, Fast and furious: The neutrophil and its armamentarium in health and disease, Med Res Rev (2023).

[9] S.P.H. Alexander, A. Christopoulos, A.P. Davenport, E. Kelly, A.A. Mathie, J.A. Peters, E.L. Veale, J.F. Armstrong, E. Faccenda, S.D. Harding, J.A. Davies, M.P. Abbracchio, G. Abraham, A. Agoulnik, W. Alexander, K. Al-Hosaini, M. Back, J.G. Baker, N.M. Barnes, R. Bathgate, J.M. Beaulieu, A.G. Beck-Sickinger, M. Behrens, K.E. Bernstein, B. Bettler, N.J.M. Birdsall, V. Blaho, F. Boulay, C. Bousquet, H. Brauner-Osborne, G. Burnstock, G. Calo, J.P. Castano, K.J. Catt, S. Ceruti, P. Chazot, N. Chiang, B. Chini, J. Chun, A. Cianciulli, O. Civelli, L.H. Clapp, R. Couture, H.M. Cox, Z. Csaba, C. Dahlgren, G. Dent, S.D. Douglas, P. Dournaud, S. Eguchi, E. Escher, E.J. Filardo, T. Fong, M. Fumagalli, R.R. Gainetdinov, M.L. Garelja, M. de Gasparo, C. Gerard, M. Gershengorn, F. Gobeil, T.L. Goodfriend, C. Goudet, L. Gratz, K.J. Gregory, A.L. Gundlach, J. Hamann, J. Hanson, R.L. Hauger, D.L. Hay, A. Heinemann, D. Herr, M.D. Hollenberg, N.D. Holliday, M. Horiuchi, D. Hoyer, L. Hunyady, A. Husain, I.J. AP, T. Inagami, K.A. Jacobson, R.T. Jensen, R. Jockers, D. Jonnalagadda, S. Karnik, K. Kaupmann, J. Kemp, C. Kennedy, Y. Kihara, T. Kitazawa, P. Kozielewicz, H.J. Kreienkamp, J.P. Kukkonen, T. Langenhan, D. Larhammar, K. Leach, D. Lecca, J.D. Lee, S.E. Leeman, J. Leprince, X.X. Li, S.J. Lolait, A. Lupp, R. Macrae, J. Maguire, D. Malfacini, J. Mazella, C.A. McArdle, S. Melmed, M.C. Michel, L.J. Miller, V. Mitolo, B. Mouillac, C.E. Muller, P.M. Murphy, J.L. Nahon, T. Ngo, X. Norel, D. Nyimanu, A.M. O’Carroll, S. Offermanns, M.A. Panaro, M. Parmentier, R.G. Pertwee, J.P. Pin, E.R. Prossnitz, M. Quinn, R. Ramachandran, M. Ray, R.K. Reinscheid, P. Rondard, G.E. Rovati, C. Ruzza, G.J. Sanger, T. Schoneberg, G. Schulte, S. Schulz, D.L. Segaloff, C.N. Serhan, K.D. Singh, C.M. Smith, L.A. Stoddart, Y. Sugimoto, R. Summers, V.P. Tan, D. Thal, W.W. Thomas, P. Timmermans, K. Tirupula, L. Toll, G. Tulipano, H. Unal, T. Unger, C. Valant, P. Vanderheyden, D. Vaudry, H. Vaudry, J.P. Vilardaga, C.S. Walker, J.M. Wang, D.T. Ward, H.J. Wester, G.B. Willars, T.L. Williams, T.M. Woodruff, C. Yao, R.D. Ye, The Concise Guide to PHARMACOLOGY 2023/24: G protein-coupled receptors, Br J Pharmacol 180 Suppl 2 (2023) S23–S144.

[10] P. Kolb, T. Kenakin, S.P.H. Alexander, M. Bermudez, L.M. Bohn, C.S. Breinholt, M. Bouvier, S.J. Hill, E. Kostenis, K.A. Martemyanov, R.R. Neubig, H.O. Onaran, S. Rajagopal, B.L. Roth, J. Selent, A.K. Shukla, M.E. Sommer, D.E. Gloriam, Community guidelines for GPCR ligand bias: IUPHAR review 32, Br J Pharmacol 179(14) (2022) 3651–3674.

[11] G.B. Downes, N. Gautam, The G protein subunit gene families, Genomics 62(3) (1999) 544–52.

[12] A.S. Hauser, C. Avet, C. Normand, A. Mancini, A. Inoue, M. Bouvier, D.E. Gloriam, Common coupling map advances GPCR-G protein selectivity, Elife 11 (2022).

[13] T. Kenakin, Bias translation: The final frontier?, Br J Pharmacol 181(9) (2024) 1345–1360.

[14] T. Kenakin, Allostery: The Good, the Bad, and the Ugly, J Pharmacol Exp Ther 388(1) (2024) 110–120.

[15] C. Dahlgren, S. Lind, J. Martensson, L. Bjorkman, Y. Wu, M. Sundqvist, H. Forsman, G protein coupled pattern recognition receptors expressed in neutrophils: Recognition, activation/modulation, signaling and receptor regulated functions, Immunol Rev 314(1) (2023) 69–92.

[16] B. Banoth, S.L. Cassel, Mitochondria in innate immune signaling, Transl Res 202 (2018) 52–68.

[17] M. Garg, S. Johri, K. Chakraborty, Immunomodulatory role of mitochondrial DAMPs: a missing link in pathology?, FEBS J 290(18) (2023) 4395–4418.

[18] B. McDonald, K. Pittman, G.B. Menezes, S.A. Hirota, I. Slaba, C.C. Waterhouse, P.L. Beck, D.A. Muruve, P. Kubes, Intravascular danger signals guide neutrophils to sites of sterile inflammation, Science 330(6002) (2010) 362–6.

[19] M. Sanchez Crespo, O. Montero, N. Fernandez, The role of PAF in immunopathology: From immediate hypersensitivity reactions to fungal defense, Biofactors 48(6) (2022) 1217–1225.

[20] K. Onnheim, K. Christenson, M. Gabl, J.C. Burbiel, C.E. Muller, T.I. Oprea, J. Bylund, C. Dahlgren, H. Forsman, A novel receptor cross-talk between the ATP receptor P2Y2 and formyl peptide receptors reactivates desensitized neutrophils to produce superoxide, Exp Cell Res 323(1) (2014) 209–217.

[21] A. Boyum, Isolation of mononuclear cells and granulocytes from human blood. Isolation of monuclear cells by one centrifugation, and of granulocytes by combining centrifugation and sedimentation at 1 g, Scand J Clin Lab Invest Suppl 97 (1968) 77–89.

[22] A. Boyum, Isolation of lymphocytes, granulocytes and macrophages, Scand J Immunol Suppl 5 (1976) 9–15.

[23] J. Bylund, H. Bjornsdottir, M. Sundqvist, A. Karlsson, C. Dahlgren, Measurement of respiratory burst products, released or retained, during activation of professional phagocytes, Methods Mol Biol 1124 (2014) 321–38.

[24] H. Lundqvist, C. Dahlgren, Isoluminol-enhanced chemiluminescence: a sensitive method to study the release of superoxide anion from human neutrophils, Free Radic Biol Med 20(6) (1996) 785–92.

[25] L. Bjorkman, H. Forsman, L. Bergqvist, C. Dahlgren, M. Sundqvist, Larixol is not an inhibitor of Galpha(i) containing G proteins and lacks effect on signaling mediated by human neutrophil expressed formyl peptide receptors, Biochem Pharmacol (2023) 115919.

[26] M. Gabl, M. Winther, A. Welin, A. Karlsson, T. Oprea, J. Bylund, C. Dahlgren, H. Forsman, P2Y2 receptor signaling in neutrophils is regulated from inside by a novel cytoskeleton-dependent mechanism, Exp Cell Res 336(2) (2015) 242–52.

[27] M.E. Falzone, R. MacKinnon, The mechanism of Galpha(q) regulation of PLCbeta3-catalyzed PIP2 hydrolysis, Proc Natl Acad Sci U S A 120(48) (2023) e2315011120.

[28] M.E. Falzone, R. MacKinnon, Gbetagamma activates PIP2 hydrolysis by recruiting and orienting PLCbeta on the membrane surface, Proc Natl Acad Sci U S A 120(20) (2023) e2301121120.

[29] T. Boussetta, H. Raad, S. Bedouhene, R. Arabi Derkawi, M.A. Gougerot-Pocidalo, G. Hayem, P.M. Dang, J. El-Benna, The peptidyl-prolyl isomerase Pin1 controls GM-CSF-induced priming of NADPH oxidase in human neutrophils and priming at inflammatory sites, Int Immunopharmacol 137 (2024) 112425.

[30] I. Fujiwara, S. Takeda, T. Oda, H. Honda, A. Narita, Y. Maeda, Polymerization and depolymerization of actin with nucleotide states at filament ends, Biophys Rev 10(6) (2018) 1513–1519.

[31] J.H. Kehrl, The impact of RGS and other G-protein regulatory proteins on Galphai-mediated signaling in immunity, Biochem Pharmacol 114 (2016) 40–52.

[32] P. Morales, M.M. Scharf, M. Bermudez, A. Egyed, R. Franco, O.K. Hansen, N. Jagerovic, J. Jakubik, G.M. Keseru, D.J. Kiss, P. Kozielewicz, O. Larsen, M. Majellaro, A. Mallo-Abreu, G. Navarro, R. Prieto-Diaz, M.M. Rosenkilde, E. Sotelo, H. Stark, T. Werner, L.M. Wingler, Progress on the development of Class A GPCR-biased ligands, Br J Pharmacol (2024).

[33] T. Andersson, C. Dahlgren, T. Pozzan, O. Stendahl, P.D. Lew, Characterization of fMet-Leu-Phe receptor-mediated Ca2+ influx across the plasma membrane of human neutrophils, Mol Pharmacol 30(5) (1986) 437–43.

[34] S. Lind, K.L. Granberg, H. Forsman, C. Dahlgren, The allosterically modulated FFAR2 is transactivated by signals generated by other neutrophil GPCRs, PLoS One 18(4) (2023) e0268363.

[35] A.J. Jesaitis, G.M. Bokoch, J.O. Tolley, R.A. Allen, Lateral segregation of neutrophil chemotactic receptors into actin- and fodrin-rich plasma membrane microdomains depleted in guanyl nucleotide regulatory proteins, J Cell Biol 107(3) (1988) 921–8.

[36] A.J. Jesaitis, K.N. Klotz, Cytoskeletal regulation of chemotactic receptors: molecular complexation of N-formyl peptide receptors with G proteins and actin, Eur J Haematol 51(5) (1993) 288–93.

[37] A.J. Jesaitis, J.O. Tolley, R.A. Allen, Receptor-cytoskeleton interactions and membrane traffic may regulate chemoattractant-induced superoxide production in human granulocytes, J Biol Chem 261(29) (1986) 13662–9.

[38] K.N. Klotz, A.J. Jesaitis, The interaction of N-formyl peptide chemoattractant receptors with the membrane skeleton is energy-dependent, Cell Signal 6(8) (1994) 943–7.

[39] K.N. Klotz, K.L. Krotec, J. Gripentrog, A.J. Jesaitis, Regulatory interaction of N-formyl peptide chemoattractant receptors with the membrane skeleton in human neutrophils, J Immunol 152(2) (1994) 801–10.

[40] H. Fu, L. Bjorkman, P. Janmey, A. Karlsson, J. Karlsson, C. Movitz, C. Dahlgren, The two neutrophil members of the formylpeptide receptor family activate the NADPH-oxidase through signals that differ in sensitivity to a gelsolin derived phosphoinositide-binding peptide, BMC Cell Biol 5(1) (2004) 50.

[41] A.J. Jesaitis, J.O. Tolley, R.G. Painter, L.A. Sklar, C.G. Cochrane, Membrane-cytoskeleton interactions and the regulation of chemotactic peptide-induced activation of human granulocytes: the effects of dihydrocytochalasin B, J Cell Biochem 27(3) (1985) 241–53.

[42] S. Lind, C. Dahlgren, R. Holmdahl, P. Olofsson, H. Forsman, Functional selective FPR1 signaling in favor of an activation of the neutrophil superoxide generating NOX2 complex, J Leukoc Biol 109(6) (2021) 1105–1120.

[43] M.W. Alnouri, K.A. Roquid, R. Bonnavion, H. Cho, J. Heering, J. Kwon, Y. Jager, S. Wang, S. Gunther, N. Wettschureck, G. Geisslinger, R. Gurke, C.E. Muller, E. Proschak, S. Offermanns, SPMs exert anti-inflammatory and pro-resolving effects through positive allosteric modulation of the prostaglandin EP4 receptor, Proc Natl Acad Sci U S A 121(41) (2024) e2407130121.

